# Generation and proof-of-concept for allogeneic CD123 CAR-Delta One T (DOT) cells in Acute Myeloid Leukemia

**DOI:** 10.1101/2022.03.15.484289

**Authors:** Diego Sánchez-Martínez, Néstor Tirado, Sofia Mensurado, Alba Martínez-Moreno, Paola Romecin, Francisco Gutiérrez-Agüera, Daniel V Correia, Bruno Silva-Santos, Pablo Menéndez

## Abstract

Chimeric Antigen Receptor (CAR)-T cells have emerged as a breakthrough treatment for relapse/refractory (r/r) hematological tumors, showing impressive complete remission rates in B-cell malignancies. However, around 50% of the patients relapse before 1-year post-treatment. T-cell “fitness” is critical to prolong the persistence and activity of the adoptively transferred product. Allogeneic T cells from healthy donors are less dysfunctional or exhausted than autologous patient-derived T cells, enabling a very attractive and cost-effective “off-the-shelf” therapy option. In this context, Delta One T cells (DOTs), a recently described cellular product based on MHC/HLA-independent Vδ1^+^ γδ T cells generated from the peripheral blood of healthy donors, represent a robust platform of allogeneic effector T cells. Here we generated and pre-clinically validated 4-1BB-based CAR-DOTs directed against the IL-3α chain receptor (CD123), a target antigen widely expressed on acute myeloid leukemia (AML) blasts. CD123CAR-DOTs showed vigorous, superior to control DOTs, cytotoxicity against AML cell lines and primary samples both *in vitro* and *in vivo*. Continuous administration of IL-15 supported the long-term persistence of a single-dose CD123CAR-DOTs in patient-derived xenograft models, sustaining their anti-leukemic efficacy as demonstrated in a re-challenge assay *in vivo*. Our results provide proof-of-concept for an allogeneic next-generation therapy based on CD123CAR-DOTs for r/r AML patients.

**KEY POINTS:**

- CD123CAR-DOTs exert specific and robust cytotoxicity *in vitro* and anti-leukemic activity *in vivo* against AML cell lines and primary cells.
- CD123CAR-DOTs show IL-15-dependent long-term persistence *in vivo* and vigorous anti-leukemic activity upon re-challenge.

## INTRODUCTION

Immunotherapy has promoted marked improvements in cancer treatment over the last decade. The immune system offers a wide range of alternatives, including cytotoxic lymphocytes – native or engineered – to eliminate treatment-resistant tumors. In this context, CD19-directed chimeric antigen receptor (CAR)-transduced T cell (CAR-T) therapies have shown impressive rates of complete remissions (CR) in relapse/refractory (r/r) B-cell malignancies, especially in B-cell acute lymphoblastic leukemia (ALL). However, one-year progression-free survival remains ~50% due to frequent relapses^3,4^. Antigen loss and phenotypic escape as well as CAR-T cell disappearance, dysfunctionality or exhaustion are commonly responsible for such failures^5,6^. Thus, new therapeutic options based on “fitter” effector T cells are being actively investigated in order to increase T-cell persistence and efficacy.

Allogeneic cytotoxic cells obtained from healthy donors (HD) are an especially attractive avenue for “off-the-shelf” next-generation CAR-T cell therapies. In particular, MHC/HLA-independent γδ T cells have emerged as a promising candidate based on their potent cytotoxic activity and release of cytokines stimulating and recruiting other immune cells to the tumor site^7–9^. While the low abundance of γδ T cells in the human peripheral blood (PB) has hindered their clinical application, recent advances in protocol development have enabled their *ex vivo* expansion to large numbers^10–13^. In particular, we have characterized Delta One T (DOT) cells, a Vδ1 T cell-enriched cellular product expressing enhanced levels of natural cytotoxicity receptors (NCRs), and demonstrated that they constitute a safe and efficient effector platform to eliminate cancer cells in *in vitro* and *in vivo* pre-clinical models of solid and hematological malignancies^10^, most notably acute myeloid leukemia (AML)^14^.

AML, the most common acute leukemia in adults, is characterized by the accumulation of differentiation-defective immature and proliferative myeloid blasts in the bone marrow (BM) and PB^15^. Unfortunately, patient overall survival has not improved significantly over the last decades, with relapses being frequent and presenting poor outcome, thus requiring hematopoietic stem cell transplantation (HSCT) as rescue therapy^16^. Despite its molecular enormous heterogeneity^15,17,18^, CD123, the α chain of the interleukin (IL)-3 receptor, is consistently expressed in >90% of AML blasts, in similar levels to leukemic stem cells, and its expression is maintained at relapse^19,20^. Based on this rational, we previously showed robust efficacy of second generation (4-1BB-based) CD123-directed CAR-T cells in preclinical AML models^21^.

Clinical implemented CAR-T cell approaches are based on abundant αβ T cells^22–24^ which have a limited applicability to the allogeneic setting due to their MHC/HLA restriction and graft-versus-host potential^25^. Importantly, an immunological BM dysfunction and T-cell exhaustion has been reported in intensively chemotherapy-treated r/r AMLs^26^. Thus, safety allowing, allogeneic T cells from HDs would represent an “off-the-shelf” cost-efficient therapy option to eliminate r/r AML. Building on these foundations, we prompted to explore the potential of MHC/HLA-independent DOT cells as vehicle for our second generation CD123 CAR in preclinical AML models.

Here we describe, for the first time, the generation of CAR-expressing DOT cells (CAR-DOTs), and provide the proof-of-concept for their application in AML treatment. Retrovirally-transfected CD123CAR-DOTs potently eliminated AML cell lines and primary samples both *in vitro* and *in vivo*, increasing the efficacy of “naked” (mock-transduced) DOTs in different models and conditions, without phenotypic alterations. Moreover, CD123CAR-DOTs released type 1 (anti-tumor) cytokines and chemokines upon exposure to primary AML cells. In *in vivo* rechallenge experiments, CD123CAR-DOTs persisted and remained functional against AML, nine weeks after their infusion into mice. IL-15 significantly improved their *in vivo* efficacy by sustaining longer action and enabling one-single-dose treatment. Our data demonstrate the potential of CAR-DOTs to represent a disruptive “off-the-shelf” allogeneic cellular immunotherapy for AML.

## METHODS

### DOT cell generation

PB mononuclear cells (PBMCs) were isolated from buffy coats from HDs by Ficoll-Hypaque gradient centrifugation. Buffy coats were obtained from the Barcelona Blood and Tissue Bank (BST) upon IRB-approval (HCB/2018/0030). PBMCs were incubated with anti-αβTCR Biotin mAb (Miltenyi Biotec) followed by incubation with anti-Biotin mAb microbeads to deplete αβT-cells by magnetic separation using AUTOMACS under depleteS protocol (Miltenyi Biotec, Biergisch Gladbach, Germany). DOT cells were generated from αβ-depleted PBMCs cultured in either 96-well plates or G-REX platform (Wilson Wolf Manufacturing), based on an adaptation from our previous protocol^10,14^. Briefly, αβ-depleted PBMCs were resuspended in OpTmizer-CTS medium supplemented with 2.5% heat-inactivated human plasma (LifeSciences), 2 mmol/L L-glutamine (Thermo Fisher) and 50 U/mL/50 ug/mL of Penicillin/Streptomycin (Thermo Fisher) and cultured for 16-20 days. Animal-free human cytokines rIL-4 (100 ng/mL), rIFNγ (70 ng/mL), rIL-21 (7 ng/mL), and rIL1β (15 ng/mL; all from PeproTech), and a soluble mAb anti-CD3 (clone OKT-3, 140 ng/mL; BioLegend), were added to the medium at day 0. At day 7, cultures were supplemented with anti-CD3 (clone OKT-3, 1 μg/mL or 2μg/mL for 96-well plate protocol), rIL-21 (13 ng/mL) and rIL-15 (70 ng/mL; also from PrepoTech). On day 11, new medium was added to cultures, supplemented with anti-CD3 (1 μg/mL) and rIL-15 (100 ng/mL). Cells were incubated at 37 °C and 5% CO_2_. DOT cells were harvested at the end of the culture and either used fresh (for *in vitro* assays) of cryopreserved (for *in vivo* experiments) in OpTmizer media + 20% human plasma + 10% DMSO and stored in liquid nitrogen.

### CD123-CAR retroviral production and transduction

To construct the retroviral transfer vector, the complete CD123-directed CAR (including CSL362 scFv, CD8 transmembrane domain, 4-1BB costimulatory domain, CD3z endodomain, and a T2A-eGFP cassette) previously described^21^ was cloned into an SFG retroviral backbone. The SFG vector expressing eGFP alone (mock vector) was used as a control. Viral particles pseudotyped with RD114 were generated using 293T cells with GeneJuice^®^ transfection reagent (Sigma-Aldrich, Saint Louis, MO, USA) following manufacturer’s instructions with a 1.5:1.5:1 μg ratio of the SFG:Peq-Pam:RD114 DNA plasmids and concentrated using Retro-X™ Concentrator (Takara, Kusatsu, Japan) following manufacturer’s instructions. DOT cells were expanded for 7 days prior to viral transduction. Proper CAR transduction was checked by flow cytometry by eGFP expression and by using AffiniPure F(ab’)_2_ Fragment Goat Anti-Human IgG (H+L) (Jackson ImmunoResearch Laboratories).

Other conditions tested during transduction optimization of DOT cells included the use of lentiviral particles as previously reported^27^ at multiplicities of infection between 10 and 50, as well as with retroviral particles with RetroNectin® (Takara) following manufacturer’s instructions. All lentiviral transduction conditions were tested at days 7 and 11 of the DOT cell expansion protocol.

### Immunophenotyping of DOT cells, cell lines and primary AML samples

Proper differentiation and activation of DOT cells was confirmed by surface staining with CD3-PB (UCHT1), CD3-APCCy7 (UCHT1), TCRγδ-PE (B1), TCRαβ-APC (IP26), CD4-APCCy7 (RPA-T4), CD8-BV510 (RPA-T8), CD25-APCCy7 (BC96), CD69-BV510 (FN50), NKp30-BV421 (P30-15), NKG2D-BV510 (1D11), CD45RA-APC (HI100), CD27-PB (O323), CD62L-BV510 (DREG-56), DNAM-1-BV510 (11A8) from BioLegend (San Diego, CA, USA) and TCR Vδ1-PE (REA173), TCR Vδ2-APC (123R3) and NKp44-APC (2.29) from Miltenyi Biotec. Briefly, 2.5×10^5^ cells were incubated with the antibodies for 30 min at 4 °C and then washed. For intracellular staining, DOT cells were incubated after membrane labelling with anti-perforin-BV241 (dG9) and anti-granzyme B-BV510 (GB11) from BD Biosciences, and anti-granzyme A-APC (CB9) from BioLegend (San Diego, CA, USA), using Fix&Perm Sample Kit (Nordic MUbio) under manufacturer’s instructions. Nonreactive, isotype-matched fluorochrome-conjugated mAbs were systematically used to set the gates. Dead cells were discarded by 7-AAD staining.

The immunophenotyping of AML cell lines and primary samples (obtained from Hospital Clínic of Barcelona) was done by surface staining with CD45-PE (HI30), CD33-BV421 (HIM3-4) and CD123-APC (7G3) (BD Biosciences, Franklin Lakes, NJ, USA). A FACSCanto™-II flow cytometer equipped with FACSDiva™ software (BD Biosciences) was used for the analysis^27^.

### *In vitro* cytotoxicity assays and cytokine release determination

The cell lines MOLM13, THP-1 and Jurkat were purchased from DSMZ (Germany) and expanded according to DSMZ recommendations. Target cells (cell lines and primary AML blasts) were labeled with 3 μM eFluor 670 (eBioscience) and incubated with CAR-DOTs or mock-DOTs at different Effector:Target (E:T) ratios for the indicated time periods in complete DOT media. CAR-DOT-mediated cytotoxicity was determined by analyzing the residual alive (7-AAD^-^) eFluor 670^+^ target cells at each time point and E:T ratio. Absolute cell counts were determined using Trucount absolute count beads (BDBiosciences). Cytokine production was measured by G-Series Human Cytokine Antibody Array 4000 from Raybiotech in supernatants harvested after 48h-incubation CAR-DOT/mock-DOT (4 donors) with 2 primary AML samples.

### *In vivo* AML patient-derived xenograft (PDX) models

7-to 12-week-old nonobese diabetic (NOD).Cg-Prkdcscid Il2rgtm1Wjl/SzJ (NSG) mice (Jackson Laboratory) were bred and housed under pathogen-free conditions in the animal facility of the Barcelona Biomedical Research Park (PRBB). Mice were intravenously (i.v.) transplanted in the tail with 2.5×10^5^ Luc-GFP-expressing primary CD123^+^ AML blasts (primograft-expanded) (579AML-PDX^LUC^)^29^. CAR- or mock-DOTs were thawed and resuspended in complete media and mice were i.v. infused in the tail vein with 10×10^6^ cells once, twice or three times 7-, 14- and 21-days post-tumor inoculation, respectively. Where indicated, mice were injected intraperitoneally with 1μg of hrIL15 every 3-4 days or daily from day 7 to the end of the experiment, as a strategy to support DOT-cell persistence *in vivo*. Tumor burden was followed by bioluminescence (BLI) using the Xenogen IVIS 50 Imaging System (Perkin Elmer). To measure luminescence, mice received 150 mg/kg of D-luciferin intraperitoneally, and tumor burden was monitored at the indicated time points as we previously described^27,30^. Living Image software (Perkin Elmer) was used to visualize and calculate total luminescence. Besides, tumor burden was followed-up at different timepoints by bleeding and BM aspirate using FACS analysis. Mice were sacrificed when control and/or mock-DOT-treated animals were leukemic, and tumor burden (hHLA-ABC^+^hCD45^+^hCD33^+^hCD123^+^ graft) and effector DOT persistence (hHLA-ABC^+^hCD45^+^hCD3^+^Vδ1^+^) were analysed in BM and PB by FACS. In re-challenge experiments, leukemia-free animals were re-infused with 2.5×10^5^ CD123+ AML primary cells, and disease reappearance was followed-up by BLI and FACS as above. All procedures were performed in compliance with the institutional animal care committee of the PRBB (DAAM9624).

### Statistical analysis

Data from at least three individual donors are shown in all figures. All *p*-values were calculated by unpaired two-tailed or one-tailed Student’s *t*-test using Prism software (GraphPad). A *p*-value *<0.05/**<0.01/***<0.001/****<0.0001 was considered statistically significant.

### Data Sharing Statement

For original data, please contact

## RESULTS

### CD123-directed DOT cells (CD123CAR-DOTs) expand robustly while preserving DOT phenotype

We set out to generate, for the first time, CAR-DOT cells, and purpose them for AML targeting. DOT cells were expanded and activated as described^10,14^, and transduction was tested at days 7 and 11 using different retroviral (PeqPam/RD114) and lentiviral (VSV-G/psPAX2) conditions (**Fig S1**). Retro-X Concentrator at day 7 showed the highest rate of anti-CD123 CAR expression and was chosen for downstream generation of CD123CAR-DOTs (**Fig 1A**). CD123CAR transduction/expression did not impact γδ TCR^+^ and Vδ1^+^ T cell percentages, which were ~97% and ~70% respectively, as in the original “naked” DOT product^10,14^. Likewise, total and Vδ1^+^ T-cell numbers at the end of the process were similar for mock- and CAR-DOTs (Fig 1C). Retroviral particles infected the Vδ1^+^ and Vδ1^-^ γδ T-cell populations in similar proportions, and the CD123CAR expression as determined by GFP positivity correlated well with anti-ScFv labelling (**Fig. 1B, C**). CAR^+^ DOTs (n=14) presented a similar T-cell differentiation stage to CAR^-^ DOTs, as demonstrated by CD62L/CD45RA staining by FACS. The majority of DOTs were effector T cells (T_EFF_), with an expected degree of variability between donors, followed by effector memory (T_EM_) DOTs, and lower contributions of central memory (T_CM_) and naïve/stem central memory (T_N/SCM_) DOTs (**Fig. 1D**). Furthermore, both CAR^+^ and CAR^-^ DOTs expressed their NK cell receptor signature according to the published^11^ hierarchy: NKG2D (>90%) > DNAM-1 (~60%) > NKp30 and NKp44. By contrast, the immune checkpoint PD-1 was strikingly absent from both DOT populations (**Fig. 1E**). The expression of the key components of cytotoxic granules, perforin (Perf), granzyme (Gzm) A and GzmB, was also not affected by CAR transduction (**Fig 1F**). Collectively, these data demonstrate the feasibility of CAR-DOT generation, and further show that retroviral infection and CD123CAR expression on the cell surface does not modify DOT expansion properties or phenotype, thus generating NK-like cytotoxic effector γδ T cells highly enriched in Vδ1^+^ T cells.

**Figure 1.**
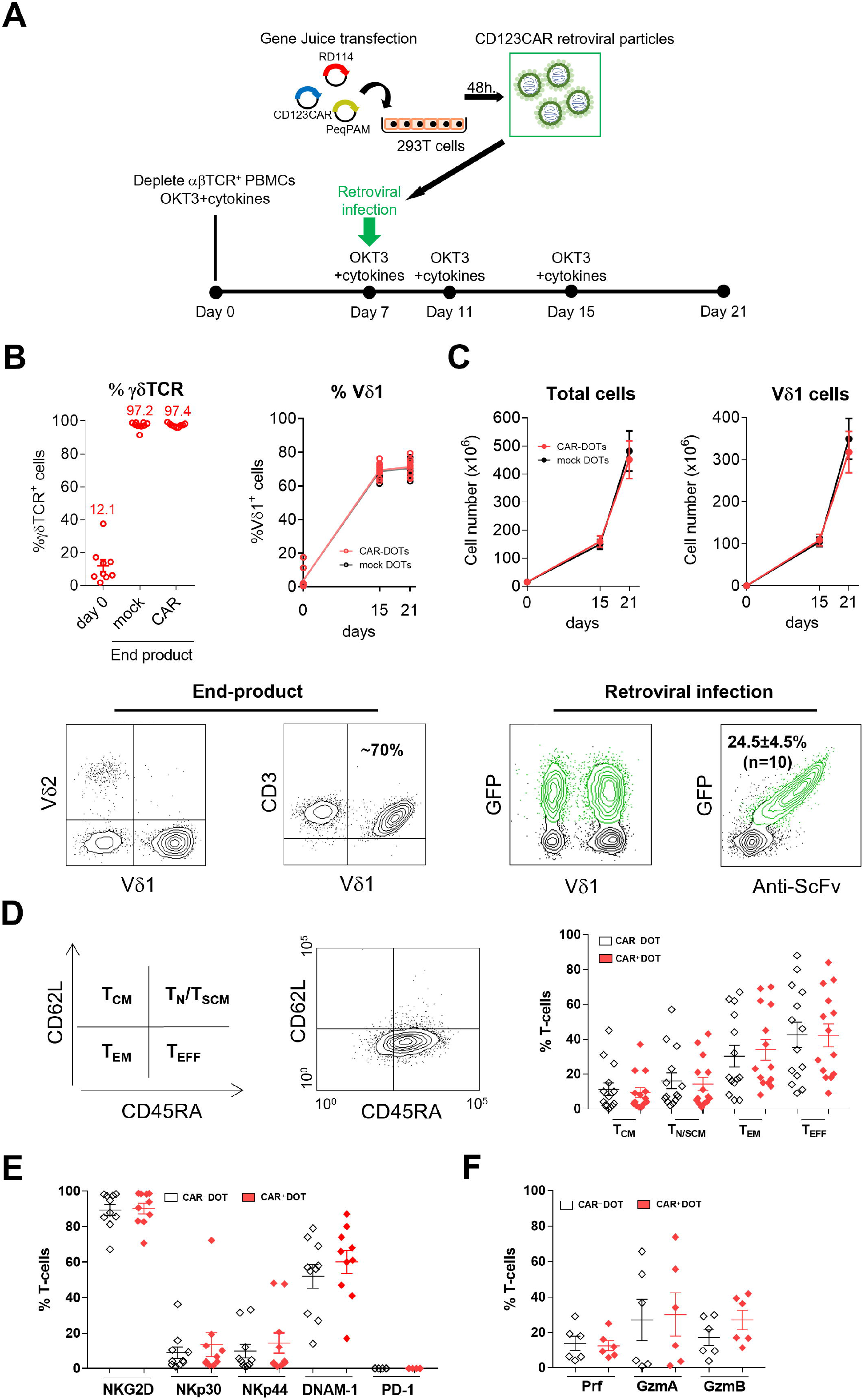
CD123CAR-DOT cell generation and phenotypic characterization. **(A) CD123CAR**-DOT cell production scheme. αβ-depleted PBMCs were cultured under OKT3 antibody and cytokine stimulation for 3 weeks and transduced with γ-retroviruses at day 7. **(B)** *Upper panels*, γδ-TCR^+^ and Vδ1 ^+^ T cell percentages evolution from day 0 to end-product for mock DOTs and CAR-DOTs. *Lower panels*, representative FACS dot plots depicting the expression of Vδ1/Vδ2 and Vδ1/CD3 in end-products. **(C)** *Upper panels*, cell number for total cells and Vδ1^+^ T cells along the CAR-DOT generation process. *Lower panels*, representative FACS dot plot for Vδ1/GFP and anti-ScFv/GFP expression. **(D)** Left-middle panels, scheme and representative FACS plot depicting T_CM_, T_N/SCM_, T_EM_, T_EFF_ T cell subsets as defined by CD62L *versus* CD45RA staining. Right panel, percentage of T_CM_, T_N/SCM_, T_EM_, T_EFF_ T-cell subsets for CAR^+^ DOT and CAR^-^ DOT subpopulations (n=14). **(E)** NKG2D, NKp30, NKp44, DNAM-1 and PD-1 expression (n=10), and **(F)** perforin (Prf), granzyme A (GzmA) and granzyme B (GzmB) expression (n=6) in CAR^-^ versus CAR^+^ DOT cells.

### CD123CAR-DOTs specifically augment cytotoxicity against AML cell lines and primary blasts *in vitro*

In order to test the anti-AML activity of CAR-DOTs, we performed *in vitro* cytotoxicity assays against the CD123^+^ AML cell lines MOLM13 and THP-1, and the CD123^-^ T-cell acute lymphoblastic leukemia line, Jurkat, as negative control. CD123CAR-DOTs specifically eliminated CD123^+^ AML cells in an E:T ratio-dependent manner, with substantially increased cytotoxic activity compared to mock-DOTs. Even at low E:T ratios (1:16, 1:8, 1:4) we achieved 20-60% more AML cell lysis than with mock-DOTs in 48h and 24h assays (**Fig 2A, S2**). Next, to evaluate their capacity to kill primary tumors, CD123CAR-DOTs were co-cultured with primary CD123^+^ AML samples, with different percentages of CD123^+^CD33^+^ blasts (**Fig 2B**). CD123CAR-DOTs showed enhanced cytotoxicity over mock-DOTs against CD123^+^ AML primary blasts in 48h assays (**Fig2C**). Moreover, compared to mock-DOTs, CD123CAR-DOTs produced significantly higher levels of master anti-tumor mediators, namely the cytokines TNF-α and IFN-γ, as well as IL-13, recently implicated in orchestrating tumor surveillance in mice^31^; the T-cell costimulator 4-1BB, a major determinant of γδ T cell-mediated tissue surveillance^32^; the signal transducer Axl, shown to maximize IL-15R signaling in human NK cell differentiation^33^; the myeloid differentiation factor, GM-CSF; and the chemokines MIP-1a (CCL3), CCL5, CXCL13 and lymphotactin (XCL1), all important in mobilizing multiple leukocyte subsets in cancer immunity^34^. While not significantly augmented, IL-2 and FasL, key mediators of T cell proliferation and cytotoxicity, respectively, also showed a tendency for higher expression in CD123CAR-DOT cells (**Fig2D**). These data demonstrate an enhanced effector potential of CD123CAR-DOT cells in direct comparison with naked/mock-DOT cells.

**Figure 2.**
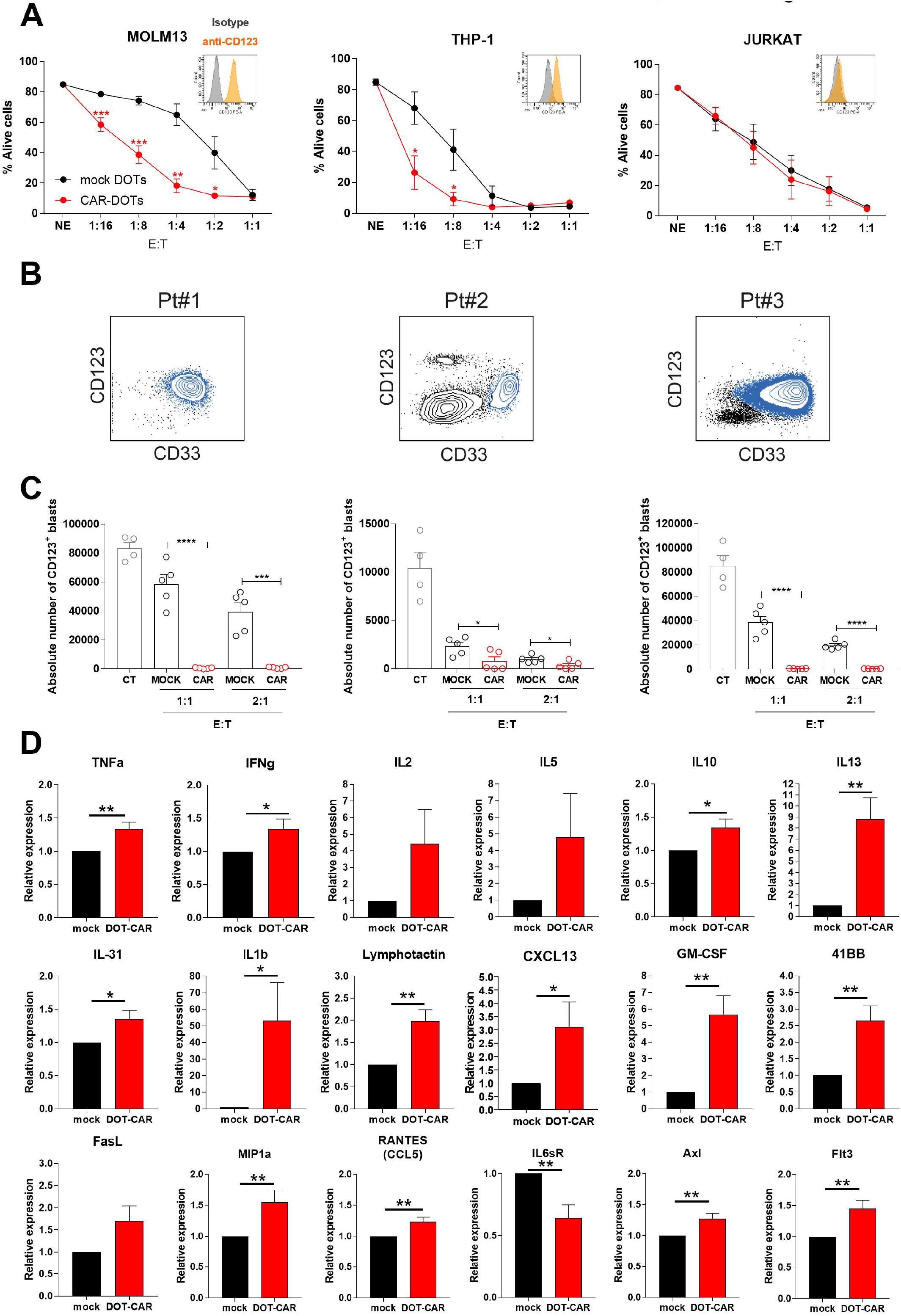
CD123CAR-DOT cells specifically target and eliminate CD123^+^ AML cell lines and primary samples in *vitro*. **(A)** Cytotoxicity of CAR-DOTs and mock-DOTs against CD123^+^ AML (MOLM13 and THP-1) and CD123^-^ T-ALL (Jurkat, negative control) cell lines at the indicated E:T ratios in 48h assays (n=5). *Small insets*, CD123 expression in each cell line. **(B)** AML primary sample (Pt, patient) phenotype showing CD33/CD123 analysis by FACS. AML blasts are highlighted in blue. **(C)** Absolute counts of alive eFluor^+^CD123^+^ AML blasts measured by FACS in 48-hour cytotoxicity assays at 1:1/2:1 E:T ratios. **(D)** Cytokine array determination in supernatants obtained from 4 donors of CAR-DOTs/mock-DOTs exposed to 2 different AML primary samples for 48h. Relative expression was normalized to mock-DOT levels. *p<0.05, **p<0.01, ***p<0.001.

### Serial infusions of CD123CAR-DOTs exhibit robust anti-leukemic effect *in vivo*

CD123CAR-DOT function was evaluated *in vivo* by employing Luc-expressing AML patient-derived xenografts (AML-PDX^LUC^). NSG mice were transplanted with 2.5×10^5^ AML-PDX^LUC^ cells and, based on our previous extensive experience with DOTs in AML xenograft models^14^, infused with 10×10^6^ (CAR- or mock-) DOTs in 3 serial infusions (one injection per week (day 7, 14, 20), starting one week post-tumor injection), and leukemia progression was followed by BLI (**Fig 3A**). CD123CAR-DOTs controlled AML tumor burden between days 20 and 47 significantly better than mock-DOTs which only managed to delay tumor growth and could not prevent an eventual logarithmic tumor growth from day 47 onwards (**Fig 3B**). At this point, we decided to provide an additional 4^th^ infusion at day 53, which resulted in complete control of leukemia in CD123CAR-DOT mice, in stark contrast with continued tumor growth in mock DOT-treated mice (**Fig. 3B**). Furthermore, FACS analyses of PB and BM samples collected at days 47 and 67 confirmed the BLI data. Leukemic burden, as determined by HLA-ABC/CD45/CD123/CD33 expression, showed high, intermediate and absent blast cells in untreated, mock-DOT and CAR-DOT-treated mice, respectively (**Fig. 3C**). These data highlight the therapeutic advantage over 12 weeks of CD123CAR-DOTs over mock-DOTs in *in vivo* AML-PDX models.

**Figure 3.**
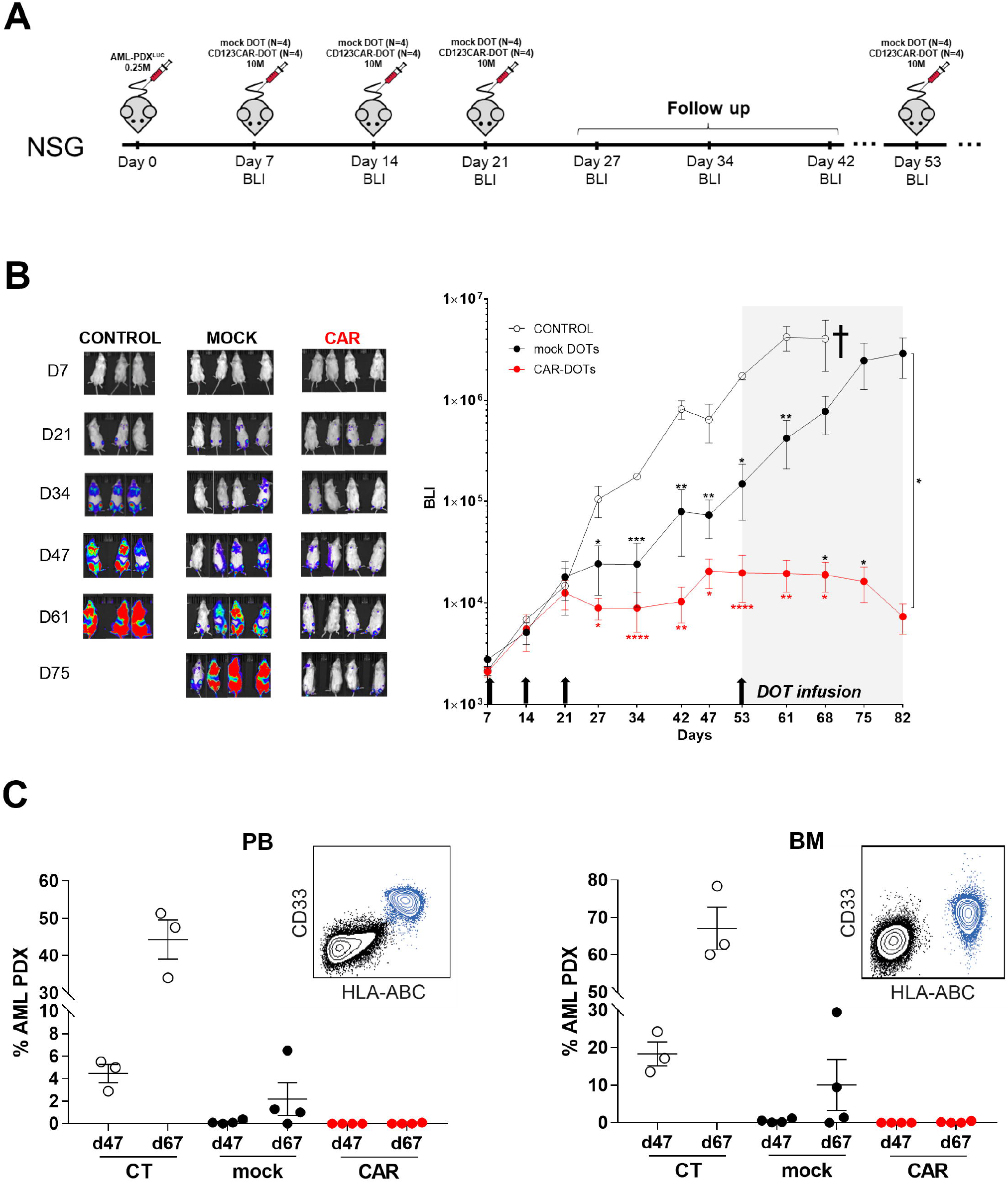
CD123CAR-DOT cells fully control growth of primary CD123^+^ AML blasts in a PDX model. **(A)** Schematic of the AML-PDX model. NSG mice (n=4/group) were i.v. injected with 2.5×10^5^ AML-PDX^LUC^ cells and then received three i.v. injections (one per week, starting 7 days after tumor injection) of 10×10^6^ CAR-DOT or mock-DOT cells. Tumor burden was monitored weekly by BLI using IVIS imaging. At day 53, mock-treated and DOT-treated mice received an extra infusion of 10×10^6^ mock-DOTs or CAR-DOTs, respectively. **(B)** IVIS imaging of tumor burden monitored by BLI at the indicated timepoints. Right panel shows the total radiance quantification (p/sec/cm^2^/sr) at the indicated timepoints. †: sacrifice. **(C)** Tumor burden analyzed by FACS in PB and BM at days 47 and 67. Primary AML blasts are shown in blue. Mouse cells are shown in black. *p<0.05, **p<0.01, ***p<0.001.

### Provision of IL-15 supports single-dose CD123CAR-DOT activity *in vivo*

Having achieved the initial proof-of-concept for CD123CAR-DOT activity in AML xenografts, we next aimed to maximize it through provision of key DOT survival factors, particularly since the murine milieu lacks human cytokines. Given that the DOT protocol relies on IL-15 as the critical cytokine in the second stage of expansion/ differentation^10,14^, we combined CD123CAR-DOTs with i.p. IL-15 administration every 3-4 days. In this set of experiments, NSG mice were transplanted with 2.5×10^5^ AML-PDX^LUC^ cells, and starting one week later, they received one, two or three doses of 10×10^6^ CD123CAR-DOTs, with or without IL-15 injection (**Fig 4A**). Leukemia engraftment was followed weekly by BLI and by flow cytometry analysis of BM and PB samples. In the absence of exogenous IL-15, three doses of CD123CAR-DOT (CAR3) were clearly more efficacious than one (CAR1) or two (CAR2) doses, since these only managed to delay tumor burden (compared to untreated control mice) (**Fig 4B,C**). Strikingly, the provision of IL-15 every 3-4 days overrode all such differences, thus allowing complete leukemia control 60 days after even with a single CD123CAR-DOT infusion (**Fig 4B,C**). These results demonstrate a critical adjuvant effect of exogenous IL-15 on CD123CAR-DOT therapy in AML-PDX *in vivo*.

**Figure 4.**
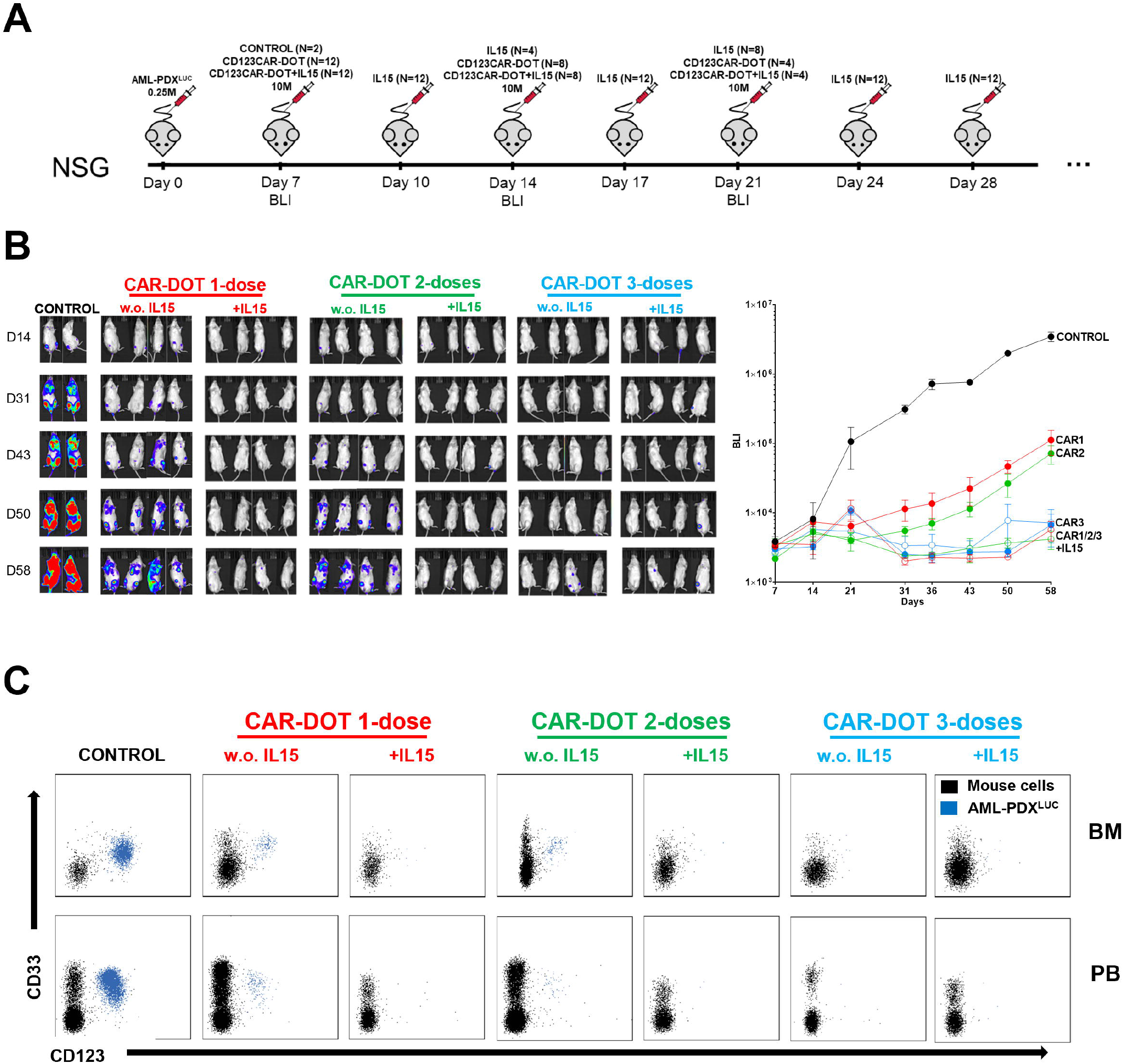
A single-dose of CD123CAR-DOT treatment combined with administration of IL-15 suffices to fully abolish AML progression in a PDX model. **(A)** Schematic of the AML-PDX model. NSG mice (n=4/group) were i.v. injected with 2.5×10^5^ AML-PDX^LUC^ cells followed 7 days after by one, two or three i.v. injections (one per week) of 10×10^6^ CAR-DOT cells, with or without administration of IL-15 i.p. **(B)** IVIS imaging of tumor burden monitored by BLI at the indicated timepoints. Right panel shows the total radiance quantification (p/sec/cm^2^/sr) at the indicated timepoints. **(C)** Tumor burden analyzed by FACS in PB and BM at the end of the experiment. AML blasts are highlighted in blue.

### Daily provision of IL-15 maximizes DOT-cell therapeutic efficacy *in vivo*

Building on the observed major impact of IL-15 on CD123CAR-DOT activity, we decided to evaluate side-by-side how IL-15 provision, in various regiments, would boost single-dose CD123CAR-DOT *versus* mock-DOT activities *in vivo*. In this experimental design, CD123CAR-DOTs or mock-DOTs were infused (in a singleone dose) one week after i.v. injection of 2.5×10^5^ AML-PDX^LUC^ cells, followed by 3 different IL-15 administration regimens: daily, every 3-4 days, or none at all (**Fig 5A**). Strikingly, mock-DOTs plus daily IL-15 showed similar leukemia control to all groups of CD123CAR-DOT-treated mice, as evaluated by BLI up to day 42 (**Fig 5B,C**). The daily provision of IL-15 was critical to enhance mock-DOT cell activity, since the 3-4-day IL-15 regimen barely reduced tumor burden compared to control groups (**Fig 5B,C**). These data were consolidated by flow cytometry analysis of BM aspirates, which showed that AML cell engraftment correlated with IVIS imaging (**Fig 5D**). Importantly, the infused (CAR- or mock-) DOTs were only detected in the BM at the endpoint of the experiment when daily IL-15 was administered, thus highlighting the importance of *in vivo* persistence of the effector T cells (**Fig 5E,F**). These data unveil the importance of in vivo IL-15 administration regimens on DOT-based therapies in AML.

**Figure 5.**
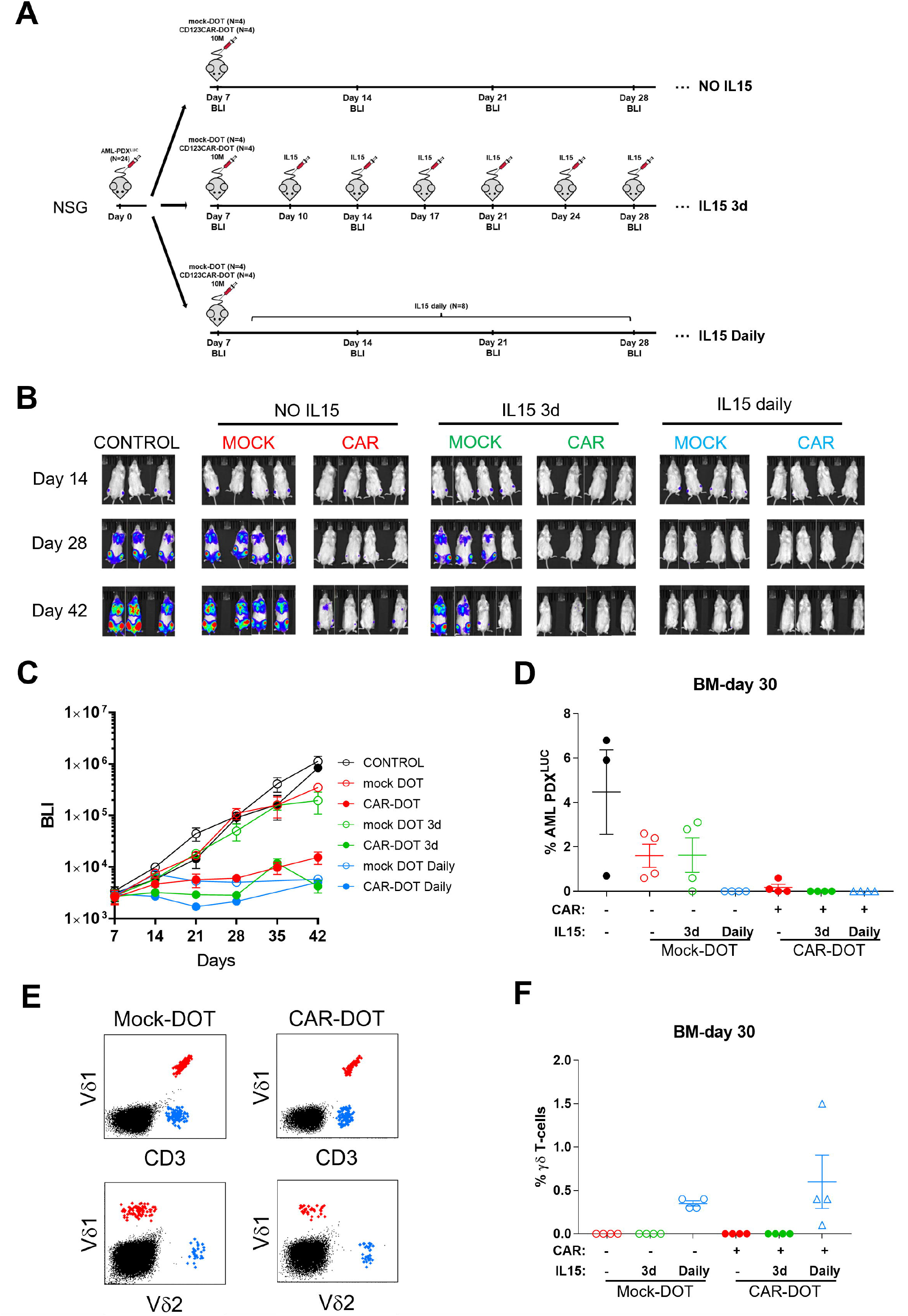
Daily-infused IL-15 increases DOT cell persistence and enables elimination of primary CD123+ AML blasts in a PDX model. **(A)** Schematic of the AML-PDX model. NSG mice (n=4/group) were i.v. injected with 2.5×10^5^ AML-PDX^LUC^ cells followed 7 days after by a one-single dose i.v. injection of 10×10^6^ CAR-DOTs or mock-DOTs. Mock- and CAR-DOT-treated mice were followed in the absence or presence of IL-15, *i.p.* administered either daily or every 3-4 days. **(B)** IVIS imaging of tumor burden monitored weekly by BLI at the indicated timepoints. **(C)** Total radiance quantification (p/sec/cm^2^/sr) at the indicated timepoints. **(D)** Tumor burden was monitored by FACS in BM at day 30. **(E)** Dot plots of representative Vδ1^+^ and Vδ2^+^ T cells determined in CAR-DOT/ mock-DOT-treated mice in BM at day 30. **(F)** Total γδ T cells were monitored by FACS in BM at day 30.

### Optimized CD123CAR-DOT treatment controls AML growth upon *in vivo* re-challenge

Persistence is a key goal and a major challenge of adoptive cellular immunotherapy. Aiming to determine the sustained functionality and potency of (CAR- or mock-) DOTs after >40 days of controlling AML progression, we then performed a tumor re-challenge experiment. Disease-free mice that eliminated the previous AML graft (**Fig 5**), either under all CD123CAR-DOT treatments or under mock-DOTs + daily IL-15, were rechallenged with an additional AML-PDX^LUC^ i.v. infusion at day 47 (**Fig 6A**). In stark contrast to the other groups, CD123CAR-DOTs plus daily IL-15 was the only condition in which tumor growth kept controlled; in fact, AML cells were barely detectable by BLI or FACS analysis of PB and BM samples (**Fig6B-E**). These results firmly demonstrate a vigorous and persistent anti-leukemia effect of CD123CAR-DOTs plus daily IL-15, sustained over 70 days and even upon tumor re-challenge, thus providing seminal proof-of-concept for their application for adoptive cellular immunotherapy in AML.

**Figure 6.**
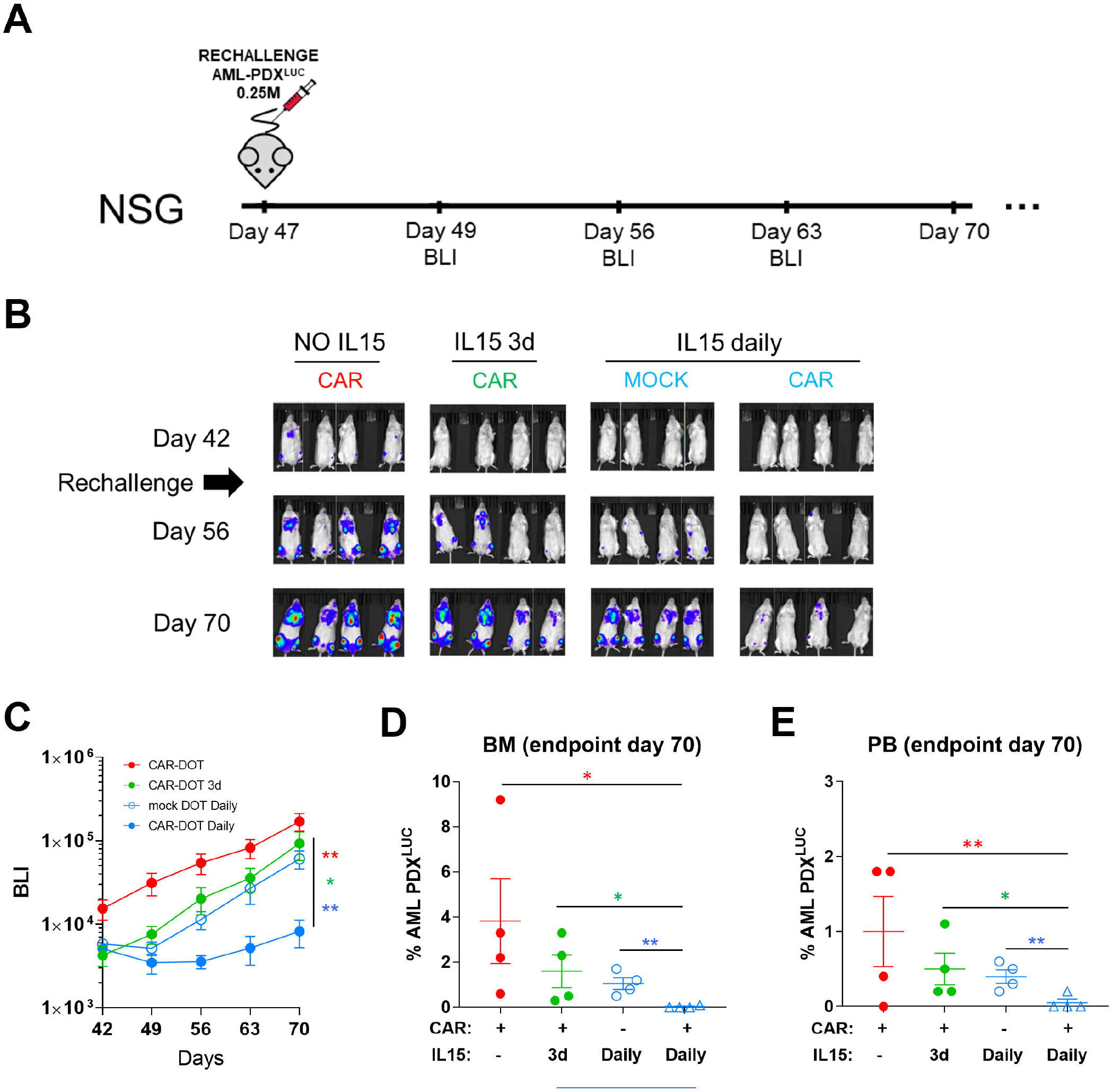
A single-dose of CD123CAR-DOT cells plus daily IL-15 sustain the ability to control AML progression upon re-challenge in a PDX model. **(A)** Schematic of the tumor re-challenge AML-PDX^LUC^ experiments. Mock-DOTs plus daily IL-15-treated mice and CAR-DOTs-treated mice (regardless of IL-15 regiment) were re-challenged with 2.5×10^5^ primary CD123+AML, 47 days after initial (CAR- or mock-) DOT cell infusion. **(B)** IVIS imaging of tumor burden monitored by BLI at the indicated timepoints. **(C)** Total radiance quantification (p/sec/cm^2^/sr) over time in mice re-challenged with AML-PDX^LUC^. Tumor burden was monitored at endpoint analysis (day 70) by FACS in BM **(D)** and PB **(E).** *p<0.05, **p<0.01, ***p<0.001 with one-tailed Student’s t-test.

## DISCUSSION

In the last few years, treatments for hematological neoplasias have undergone a revolution due to immunotherapy. In B-cell malignancies, autologous CAR-T cell therapies have rescued r/r patients on the path to palliative care, obtaining very impressive CR rates^1,2^. The number of CAR-T clinical trials keeps increasing, aiming to implement alternatives for bad prognosis tumors and expanding their horizons to other types of diseases (HIV, COVID-19, autoimmune diseases, cardiac damage, etc.)^35–37^. However, CAR T cells are still at the beginning of their clinical implementation, and a significant fraction (around 50%) of the patients relapse after treatment in a relative short period of time^3,4^. Factors like the low number of T cells retrieved from the patient’s leukapheresis, tumor blast contamination in the starting material, or T-cell exhaustion in multi-treated patients, can all explain autologous CAR-T failures stemming from the manufacturing process^38^.

The BM of leukemia patients is exposed to several cycles of chemotherapy and immunosuppressants, which alters the status of normal T cells^5,39,40^. Particularly in AML patients, T cells have been characterized with an exhausted phenotype at relapse^26^. In this context, allogeneic T cells represent an attractive alternative, since circulating T cells from HDs have not suffered disease- or therapy-induced dysfunction. While various groups have reported the generation of allogeneic αβT-cells, these cells must undergo genetic manipulation, generally CRISPR/Cas9 editing, to eliminate the endogenous T-Cell Receptor (TCR) since it would otherwise drive a (potentially severe) graft-versus-host reaction. Elegant studies have reported the feasibility of these TCR^KO^ CAR-T cells in preclinical models and even in pediatric patients^41–43^. However, a relevant recent paper has reported that TCR elimination in CAR-T cells leads to a decrease in persistence^44^. Furthermore, from a clinical point-of-view, CRISPR/Cas9 handling would still have to solve complex regulatory requirements.

Since the vast majority of γδ T cells are HLA-independent^45^, they do not require genetic manipulation to be used as allogeneic therapies. It was with this purpose that we previously developed DOT cells, a Vδ1 T cell-enriched cell product characterized by the upregulation of multiple activating/ cytotoxicity-associated NK-cell receptors (NKG2D, DNAM-1, NKp30, NKp44), and showed their anti-leukemic potential^10,14^. However, our previous data in AML xenograft models left a significant margin for improvement in terms of control of tumor burden^15^. Here we investigated whether DOT transduction with a CD123-directed CAR would improve efficacy and lead to complete and durable leukemia remissions. We show that, compared (side-by-side) with control (mock-)DOTs, CAR123-DOTs are substantially more potent at eliminating AML blasts (both cell lines and primary samples) *in vitro*; release increased levels of key cytokines involved in immune orchestration and amplification of the anti-tumor response; and are more efficient at controlling tumor growth *in vivo*, in AML-PDX experiments conducted over 70 days.

We also found that upon administration of IL-15 in our AML-PDX models, CD123CAR-DOTs (and also mock-DOTs) persisted better, thus allowing a single dose to maintain mice free of leukemia. Critically, daily-infused IL-15 enabled CD123CAR-DOTs to sustain their anti-leukemic activity for more than 70 days, even after AML re-challenge. These results are in agreement with recent elegant publications showing CAR-Vδ1 T-cell efficacy and the adjuvant effect of IL-15 in B-cell lymphoma and hepatocellular carcinoma models^12,46^. Only the clinical setting can establish the real benefit of IL-15 co-administration (potentially with an inducible “switch off” system) with CD123CAR-DOTs. Obviously, our (and other) pre-clinical mouse xenograft models lack endogenous human IL-15; thus, basal circulating IL-15 levels in patients could suffice to sustain CD123CAR-DOT persistence and activity/efficacy in the clinical setting. Importantly, lymphodepletion regimens that are used prior to CAR T-cell infusions were shown to increase the levels of IL-15 in patients^47^. Therefore, although described as safe in non-human primates^48^, the need for exogenous (adjuvant) IL-15 should be carefully examined since it caused some toxicities in a recent clinical trial in cancer patients^49^. This notwithstanding, this clinical trial employed a continuous bolus of IL-15 as single therapy, and thus, lower and/ or intermittent IL-15 dosing may be a promising solutions^50,51^.

An important final aspect to discuss is the potential advantage of using DOTs as effector cells for CAR transduction. Compared with conventional αβ-dominated T-cell products, DOTs offer several advantages: first and foremost, the previously mentioned suitability for allogeneic use, without any need for additional genetic engineering^52^; additionally, the lack of T-cell exhaustion mediated by the immune checkpoint PD-1 (absent in DOTs^11^ and CD123CAR-DOTs, see Fig. 1E), and the upregulation of a NK-associated cytotoxicity machinery (including DNAM-1, NKp30 and NKp44)^10,14^ that may be critical in the advent of CAR antigen loss as previously observed with CD19 in B-ALL^6^. Importantly, this enhanced NK-like cytotoxicity is also an advantage over other γδ T cell-based products being developed for adoptive immunotherapy of cancer^7-9,12,46^. On the other hand, in comparison with NK cells, which obviously share such activating NK-cell receptors, DOTs offer as key advantages^7,8,53^ the previously documented^11^ absence of inhibitory KIRs, namely KIR2DL1, KIR2DL2, KIR2DL4, KIR2DL5A and KIR3DL1, and the additional input of TCR-dependent activation and expansion (up to very high yields as reported^10,14^). In fact, the presence of a stochastically recombined and highly polyclonal TCR repertoire, allowing HLA-unrestricted recognition of stress-induced molecules^8^, may also counteracts tumor immune evasion (via CAR antigen loss or HLA downregulation, for example). Moreover, the “trophic” (homeostatic) signals received through γδ TCR binding to butyrophilins constitutively expressed in multiple tissues^54^ may be beneficial for *in vivo* persistence, and may underlie the recent description of a striking Vδ1^+^ T cell-associated CD19CAR-T cell expansion (from a very low initial frequency), over 10 years in a disease-free CLL patient^55^. Furthermore, although allogeneic CD19-directed CAR-NK cells have recently shown promising clinical results^56^, their “off-the-shelf” implementation may be difficult due to the reported substantial decrease in viability after the freezing/thawing process^57,58^, which contrasts with the robustness we observed with CD123CAR-DOTs. We therefore strongly believe that CAR-DOTs combine a set of properties that may revolutionize allogeneic cell immunotherapy of hematological (and possibly also solid) cancers in the near future.

## Acknowledgments

We are indebted with Dr. Talía Velasco-Hernández for technical support with IL-15 infusions, Dr. Matteo Libero Baroni for retroviral cloning, Prof. Irmela Jeremias for the Luc+ AML primografts, Dr. Maksim Mamonkin for retroviral generation protocol kindly provided, Dr. Clara Bueno and Dr. Virginia Rodríguez for lab maintenance, Dr. Pedro Roda for technical support, and Natacha Gonçalves-Sousa for administrative help. This research was funded by “la Caixa” Foundation (ID 100010434) under the agreement LCF/PR/HR19/52160011 to BS-S and PM; and also by ISCIII-RICORS within the Next Generation EU program (plan de recuperación, transformación y resilencia). PM also thanks CERCA/Generalitat de Catalunya and Fundació Josep Carreras-Obra Social la Caixa for core support. DS-M is partially supported by a Sara Borrell fellowship from the Instituto de Salud Carlos III. NT has a FPU PhD Scholarship form MINECO.

## Author contributions

DS-M conceived the study, designed, and performed experiments, analyzed data and wrote the manuscript. NT, SM, AM-M, PR, FG-A, DVC performed experiments and analyzed data.BS-S and PM conceived the study, designed experiments and wrote the manuscript.

