## Supplementary material for "Generation and proof-of-concept for allogeneic CD123 CAR-Delta One T (DOT) cells in Acute Myeloid Leukemia": Suppl. Material

**SUPPLEMENTARY FIGURES**

Sánchez-Martínez, D. et al. Figure S1

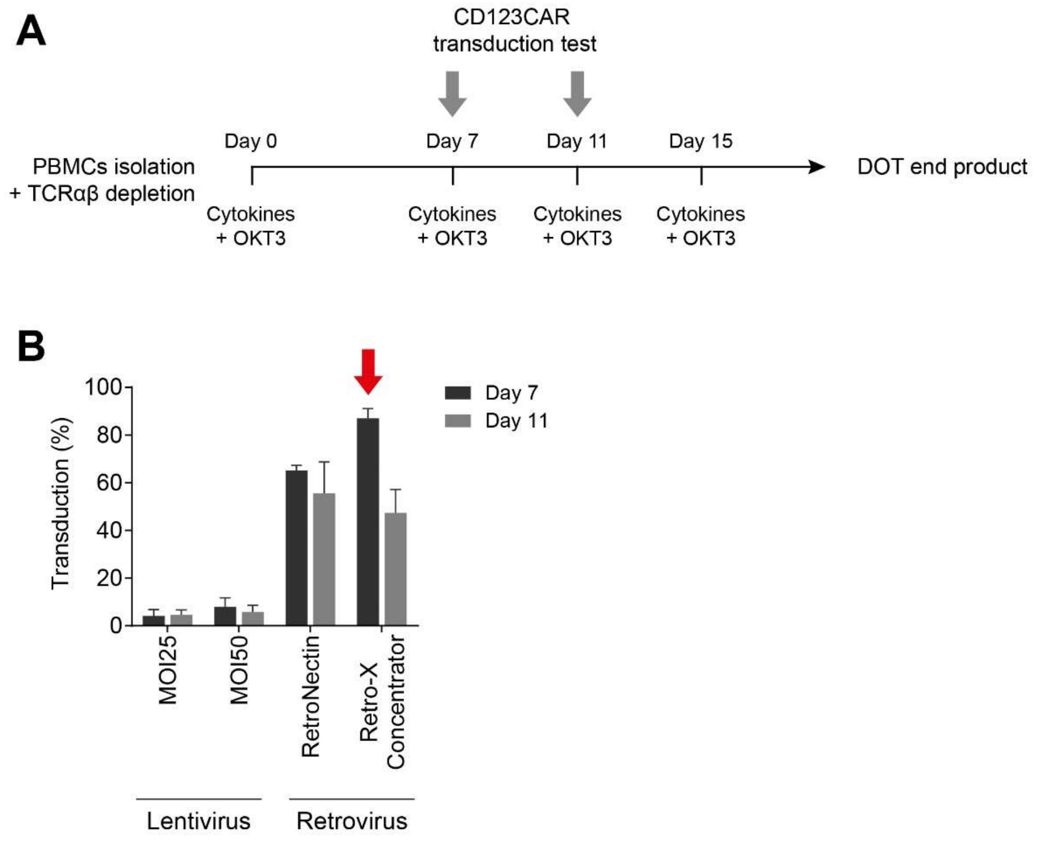


**Figure S1. CD123CAR-DOT transduction protocol establishment. (A)** CD123CAR-DOT transduction scheme. The cells were transfected at days 7 or 11 of the DOT protocol. **(B)** MOI (Multiplicity of infection) 25 and 50 for lentiviruses, RetroNectin and Retro-X Concentrator for retroviruses, were tested at day 7 or 11 to transfect DOT cells.

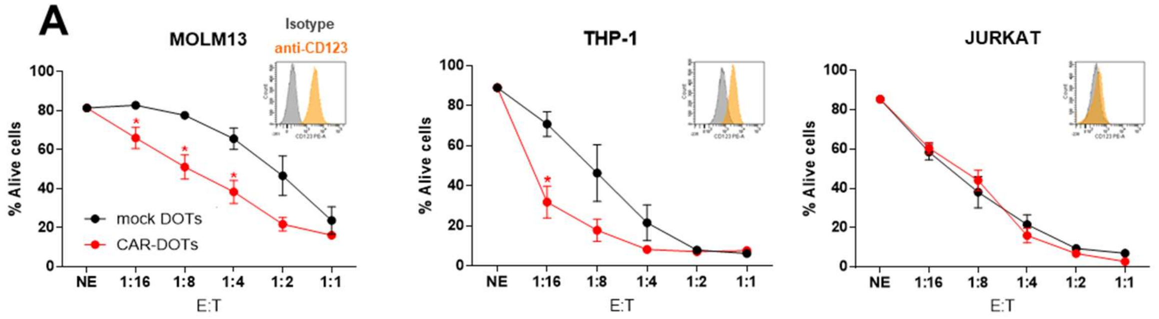


**Figure S2. CD123CAR-DOTs specifically target and eliminate CD123+ AML cell lines *in vitro*. (A)** Cytotoxicity of CAR-DOTs and mock-DOTs against $CD123^{+}$ AML (MOLM13 and THP-1) and $CD123^{-}$ T-ALL (Jurkat, negative control) cell lines at the indicated E:T ratios in 24-hour assays (n=5). *Small insets*, CD123 antigen density in each cell line. *p<0.05, **p<0.01, ***p<0.001.
